# Gut feelings: Associations of emotions and emotion regulation with the gut microbiome in women

**DOI:** 10.1101/2022.05.26.493641

**Authors:** Shanlin Ke, Anne-Josee Guimond, Shelley S. Tworoger, Tianyi Huang, Andrew T. Chan, Yang-Yu Liu, Laura D. Kubzansky

**Author notes:** Corresponding authors: Y.-Y.L. and L. D. K. These authors contributed equally (Shanlin Ke, Anne-Josee Guimond).

## Abstract

Accumulating evidence suggests that positive and negative emotions, as well as emotion regulation processes, influence the onset and the progression of multiple human diseases. However, the underlying mechanisms connecting these emotion-related factors with physical health are not fully understood. Recent work linking the gut microbiome with both mental and physical health suggests it may be a potential pathway. Yet, its association with emotions and emotion regulation are understudied. In this study, we examined whether positive and negative emotions, as well as two emotion regulation strategies (i.e., cognitive reappraisal and suppression), were associated with the diversity, overall structure, and specific species/pathways of the gut microbiome in 206 healthy women. We found that the alpha diversity was negatively associated with suppression. Moreover, using multivariate analysis, we found that positive emotions were inversely associated with the relative abundance of *Firmicutes bacterium CAG 94* and *Ruminococcaceae bacterium D16*, while negative emotions were directly related to relative abundance of these same species. Moreover, we found associations of emotions and emotion regulation with microbial metabolic pathways. For example, negative emotions were inversely related to biosynthesis of pantothenate, coenzyme A, and adenosine. Taken together, our findings offer human evidence supporting linkages of emotions and related regulatory processes with the gut microbiome. These findings highlight the critical importance of incorporating the human gut microbiome in our understanding of emotion-related factors and their associations with physical health.

## Introduction

Mounting evidence supports the role of emotion-related factors in disease etiology and health promotion^1,2^. Both negative (e.g., depression, anxiety, post-traumatic stress disorder, and negative affect) and positive (e.g., happiness, excitement, curiosity, and positive affect) manifestations of emotion have been linked with maintaining physical health as well as risk of developing disease, including cardiovascular disease (CVD)^2-5^, obesity^6^, cancer^7^, and overall mortality^8^. Additional work suggests emotion regulation – that is, the strategies by which individuals shape the nature of emotions they experience as well as when and how they experience these emotions – may also impact health and help to explain observed associations of distinct positive and negative emotion-related factors with a range of health outcomes^9-11^. While investigators hypothesize that emotion-related factors influence health through direct biological alterations or via influencing behaviors that affect biological processes,^2,12^ our understanding of specific biological mechanisms connecting emotion and emotion regulation with physical health is limited.

The human body and particularly the gastrointestinal (GI) tract is inhabited by hundreds of trillions of microbes, collectively known as the microbiome. Rather than simple passengers in or on our bodies, commensal microbes play key roles in physical health and diseases^13^. A disrupted gut microbiome is associated with various physical health conditions including cardiometabolic diseases^14^. Moreover, accumulating evidence suggests there is bidirectional communication between the resident microbes of the GI tract and the brain, i.e., the gut-brain axis, which plays a key role in maintaining brain health^15^. Recent work further suggests associations between the gut microbiota perturbations and multiple psychiatric disorders that are characterized by the experience of frequent and intense negative emotions, such as depression^16^ and anxiety^17^. Taken together, these studies suggest that certain gut microbial species or functions could be a key mechanistic pathway by which emotion-related factors contribute to physical health outcomes.

Few studies have formally tested whether gut microbial profiles respond to or modulate emotion-related factors. Recently, a small cross-sectional study based on 16S rRNA gene sequencing data within a Korean cohort (n= 83) linked positive emotions to a number of microbial features, such as *Prevotella* enterotype, alpha diversity (i.e., within-sample taxonomic diversity), and a novel genus from the family Lachnospiraceae^18^. However, 16S rRNA gene sequencing typically yields general taxonomic profiling (e.g., family or genus), potentially missing associations with specific microbial species and function.

Links between emotion regulation strategies and the gut microbiome are currently unexplored. Beyond the effects of specific emotions, the *regulation* of emotion appears to be critical for physical health^10^. Emotion regulation is a higher order feature of emotional functioning that involves both up-and down-regulation of positive and negative emotions and reflects the ability to manage environmental demands^10^. Two major types of emotion regulation strategies have been studied: response-focused strategies employed after an emotion has occurred (e.g., suppression) and antecedent-focused strategies used before the emotional response is fully deployed (e.g., cognitive reappraisal). Suppression consists of inhibiting behaviors, such as facial (e.g., repressing crying or laughing) or verbal expressions, to withhold information from others (and possibly oneself) about emotional states. Research generally suggests suppression is maladaptive for physical health with multiple short- and long-term costs (e.g., increasing the negative emotions that individuals seek to suppress, increased activation of the sympathetic nervous system)^19^. Suppression has been associated with higher inflammation levels and higher CVD risk^10^, and with greater risk of all-cause and cancer mortality^20^. In contrast, cognitive reappraisal, which involves modifying one’s assessments of a given situation either to attenuate or amplify the attendant emotional experience, appears more adaptive, with a less deleterious impact on sympathetic nervous system activation or cognitive resources^19^; it has also been associated with favorable health outcomes, including reduced risk of CVD^10^. No work has yet examined if specific profiles of gut microbiota composition and functional capacity are associated with suppression and cognitive reappraisal. Identifying plausible mechanistic pathways linking emotion-related factors and health is a necessary step towards establishing causality in emotion-health associations and understanding how interventions targeting emotional functioning can protect health.

The primary goal of this study is to examine associations of positive and negative emotions, as well as adaptive and maladaptive emotion regulation strategies (i.e., cognitive reappraisal and suppression) with the gut microbiota compositions and functional capacity. To do this, we used an agnostic approach to evaluate emotion-related factors in relation to features of the gut microbiome using data from a sample of women participating in an ongoing cohort of middle-age and older women^21^. Based on previous work suggesting that positive emotions and cognitive reappraisal are generally health-protective, and that negative emotions and suppression are associated with poorer health outcomes^2,4,9,22^, we expected to observe different associations between these two sets of processes with features of the gut microbiome. However, we did not have *a priori* hypotheses regarding the nature and the magnitude of these associations.

In this study, we evaluated associations of the gut microbiome with both positive and negative emotions, as well as emotion regulation strategies. We first evaluated associations with microbiome diversity. We then evaluated the effect of a range of potential confounders, including sociodemographics (e.g., age, marital status)^23,24^, health-related factors (e.g., history of hypertension, body mass index [BMI])^25,26^, and health behaviors (e.g., physical activity, diet)^27,28^. Finally, we assessed associations of the emotion-related factors with taxonomic profiles at the species level and with functional profiles at the metabolic pathway level.

## Results

### Characteristics of study population

Data are from 206 women participating in a sub-study (Mind Body Study: MBS^21^) conducted from 2013-2014 within a large ongoing cohort study of professional women (Nurses’ Health Study II) (Fig. 1). The MBS included a detailed psychosocial questionnaire at two time points one year apart and up to four stool collections in the interval period^21^. Briefly, emotion-related factors were self-reported and included positive and negative emotions, cognitive reappraisal, and suppression. To measure positive emotions, we used two positively-worded items (“I was happy”, “I felt hopeful about the future”) from the 10-item Center for Epidemiological Studies Depression (CESD-10) Scale^29^, with higher scores indicating higher levels of positive emotions. For negative emotions, we derived a score using four items from the CESD-10^29^ (“I was bothered by things that usually don’t bother me”, “I felt depressed”, “I felt fearful”, “I was lonely”), and three items from the Kessler Psychological Distress Scale, 6-item version^30^ (K-6; During the past month, about how often did you feel… “nervous”, “hopeless”, “restless or “fidgety”). Higher scores indicate higher levels of negative emotions. We also included the Emotion Regulation Questionnaire, a validated 10-item scale that measures cognitive reappraisal (six items) and suppression (four items)^31^, with higher subscale scores reflecting higher use of each emotion regulation strategy. All emotion-related factor scores were standardized for analyses using Z-score (Mean = 0, SD = 1). All MBS participants were invited to provide up to two pairs of stool samples, 6 months apart, approximately three months after the initial psychosocial survey and three months before the follow-up survey. Each stool sample pair was collected from two bowel movements one to three days apart using a vial with RNALater as the preservative. DNA purification from stool aliquots were performed according to standard protocols used in the Human Microbiome Project (HMP)^32,33^. We performed whole-metagenome shotgun (WMS) sequencing to get both taxonomic and functional profiles of the gut microbiome samples using MetaPhlAn3^34^ and HUMANn3^34^.

**Fig. 1.**
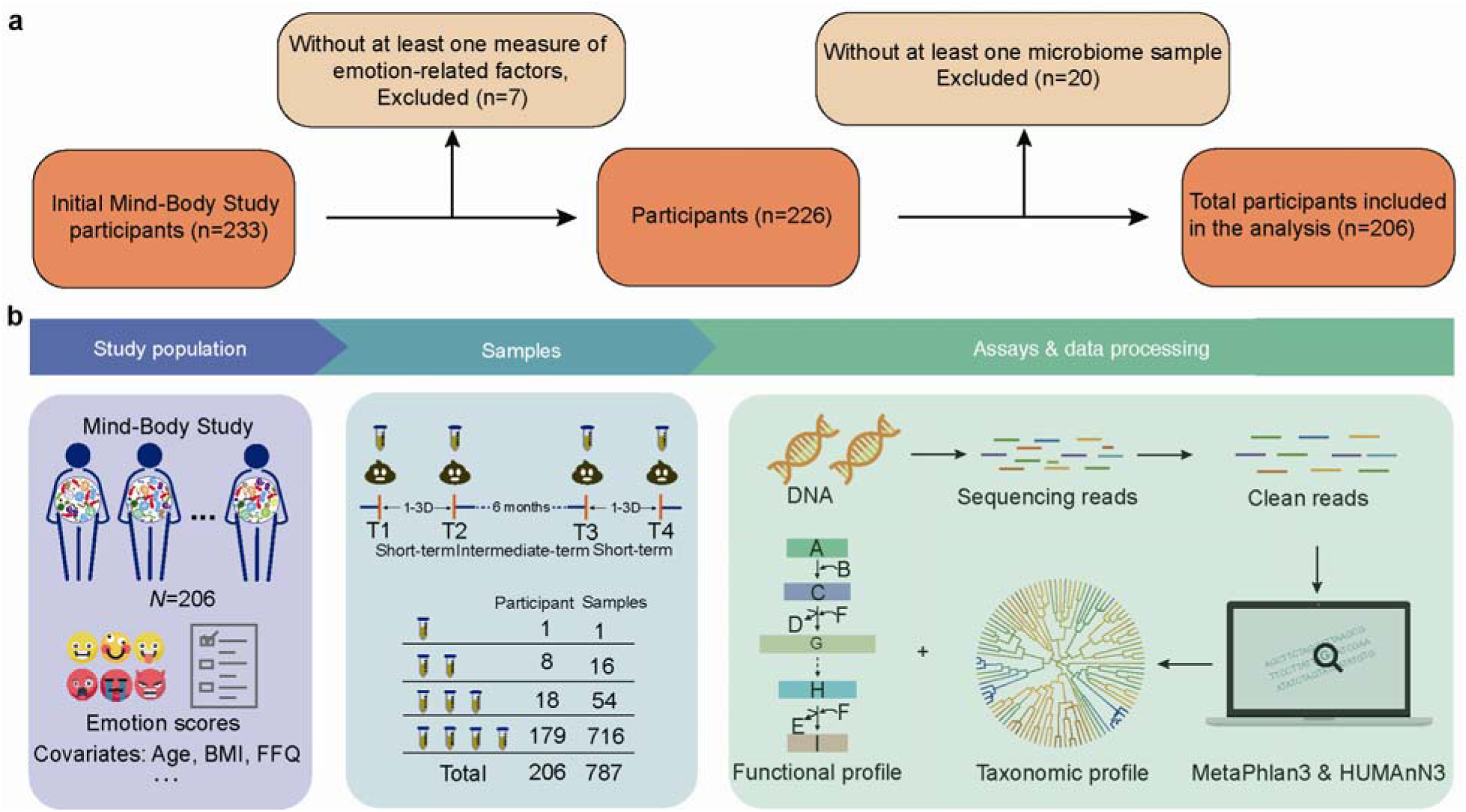
Conceptual framework of study. This analysis was designed to evaluate the associations of emotion-related factors (i.e., positive and negative emotions, as well as emotion regulation strategies), with the gut microbiome. **a**, The exclusion process for participants and the microbiome samples. **b**, The analytic sample included 206 women from the Mind Body Study, nested within the Nurses’ Health Study II cohort. Each participant provided up to four stool samples; one pair of stool samples was collected 24–72 h apart about three months after the questionnaire was administered followed by a second pair about 6 months later. Phenotypic data was collected through mailed questionnaire assessment, including individual characteristics, emotion-related factors, health conditions and health behaviors including habitual dietary intake. DNA was extracted from all fecal samples. The taxonomic and functional profiling were performed using MetaPhlAn3 and HUMAnN3, respectively. D=day; BMI=body mass index; FFQ=food frequency questionnaire.

Table 1 presents basic characteristics of the study population. Participants were on average 60.7 years old in 2013 (SD=3.8; range=49.4-66.8). Most participants were White (96%) and married/cohabitating (78%). Mean BMI was 26.4 kg/m^2^, 33% of women reported a history of hypertension, and 5% reported a history of diabetes. Higher positive emotion scores were inversely correlated with lower negative emotion and suppression scores (r = -0.49, *p* <.0001 and -0.30, *p* <.0001, respectively), and positively correlated with higher cognitive reappraisal scores (r = 0.24, *p* = .001. Higher negative emotion scores were also correlated with higher suppression scores (r = 0.28, *p* <.0001); see Supplemental Table 1.

**Table 1.**
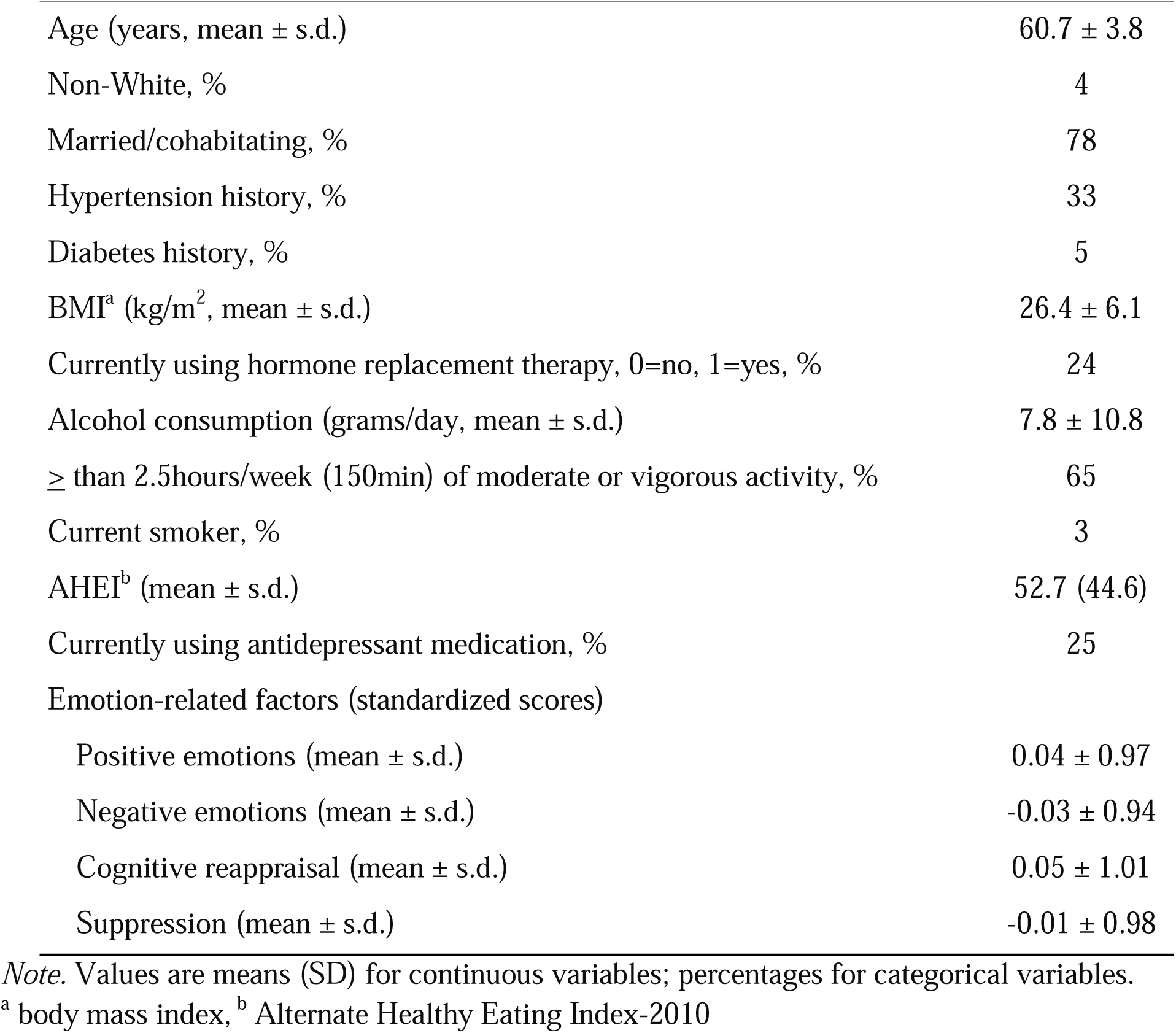
Characteristics of the study population at 2013 baseline (N=206)

### Emotion-related factors and microbiome diversity

In total, we identified 467 microbial species and 479 metabolic pathways across all microbiome samples. First, we used the Richness, Evenness, Shannon, and Simpson Diversity indices to determine the alpha diversity for the microbiome samples at the species and metabolic pathway levels. We then calculated Spearman correlations between emotion-related factors and the alpha diversity. If participants provided multiple microbiome samples (n=205), we computed the average alpha diversity across samples for each participant. We observed higher levels of suppression were significantly associated with lower Evenness and Simpson diversity at the species level (Fig.2a). However, the alpha diversity was not significantly correlated with positive emotions, negative emotions, or cognitive reappraisal. Further, we found no significant associations at the metabolic pathway level.

**Fig. 2.**
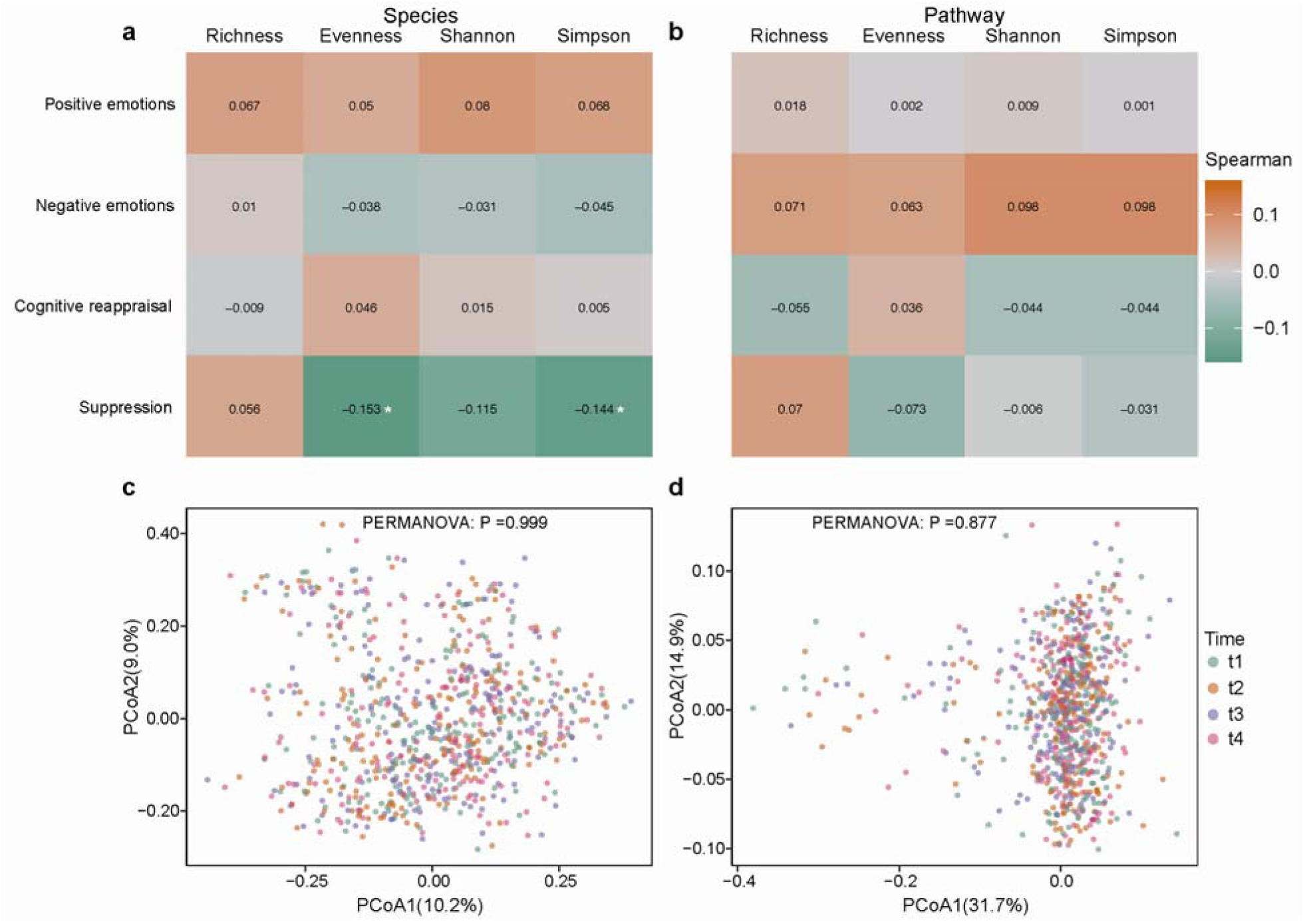
The association between emotion-related factors and gut microbial diversity. The heatmap displays the correlation between average alpha diversity and emotion-related factors at the species (**a**) and metabolic pathway (**b**) levels. Correlations were determined by Spearman correlations and asterisks denote statistically significant associations (*p* **≤** 0.05). Associations in this panel were conducted based on average alpha diversity of microbiome collected from 206 participants. Principal Coordinates Analysis (PCoA) of all microbiome samples over time at (**c**) the species and (**d**) the metabolic pathway levels based on Bray-Curtis (BC) dissimilarity. Analyses in panels **c**-**d** were conducted based on all 787 metagenomes collected from 206 participants. All PERMANOVA tests were performed with 9999 permutations based on Bray– Curtis dissimilarity.

Because the microbiome samples were collected at multiple time points, we also assessed if the microbiome features changed meaningfully over time using permutational multivariate analysis of variance (PERMANOVA, a geometric partitioning of variation across a multivariate data cloud, defined explicitly in the space of a chosen dissimilarity measure, in response to one or more factors in an analysis of variance design). Specifically, PERMANOVA (n=9999 permutations) revealed no significant differences over time with respect to both species (Fig.2c, *p*=0.999) and metabolic pathway s (Fig2.d, *p*=0.877). This is concordant with prior work with the same sampling strategy showing that the male adult gut microbiome remains relatively stable over time^35^.

### Associations between host factors, emotion-related factors, and overall gut microbiome structures

We next sought to determine if host factors as measured by sociodemographic, health-related, and health behavior covariates are significantly associated with overall microbiome structure at both taxonomic and functional levels at each time point. Using PERMANOVA analysis, considering each factor in separate models, we found three host factors (i.e., physical activity, BMI, and Type 2 diabetes history) were significantly associated with overall structure in both taxonomic and functional profiles at each time point (Fig.3). Among all host factors considered, physical activity accounted for the largest proportion of variation in the species-level taxonomic profiles (Fig.3a). It also explained the greatest amount of variance of the functional profiles at metabolic pathway level from the second and fourth time points (Fig.3b). Notably, our analysis also found that negative emotion scores were significantly associated with overall microbiome structure at the metabolic pathway level at three time points (Fig.3). Positive emotion, cognitive reappraisal, and suppression scores were not significantly associated with overall microbiome configurations.

**Fig. 3.**
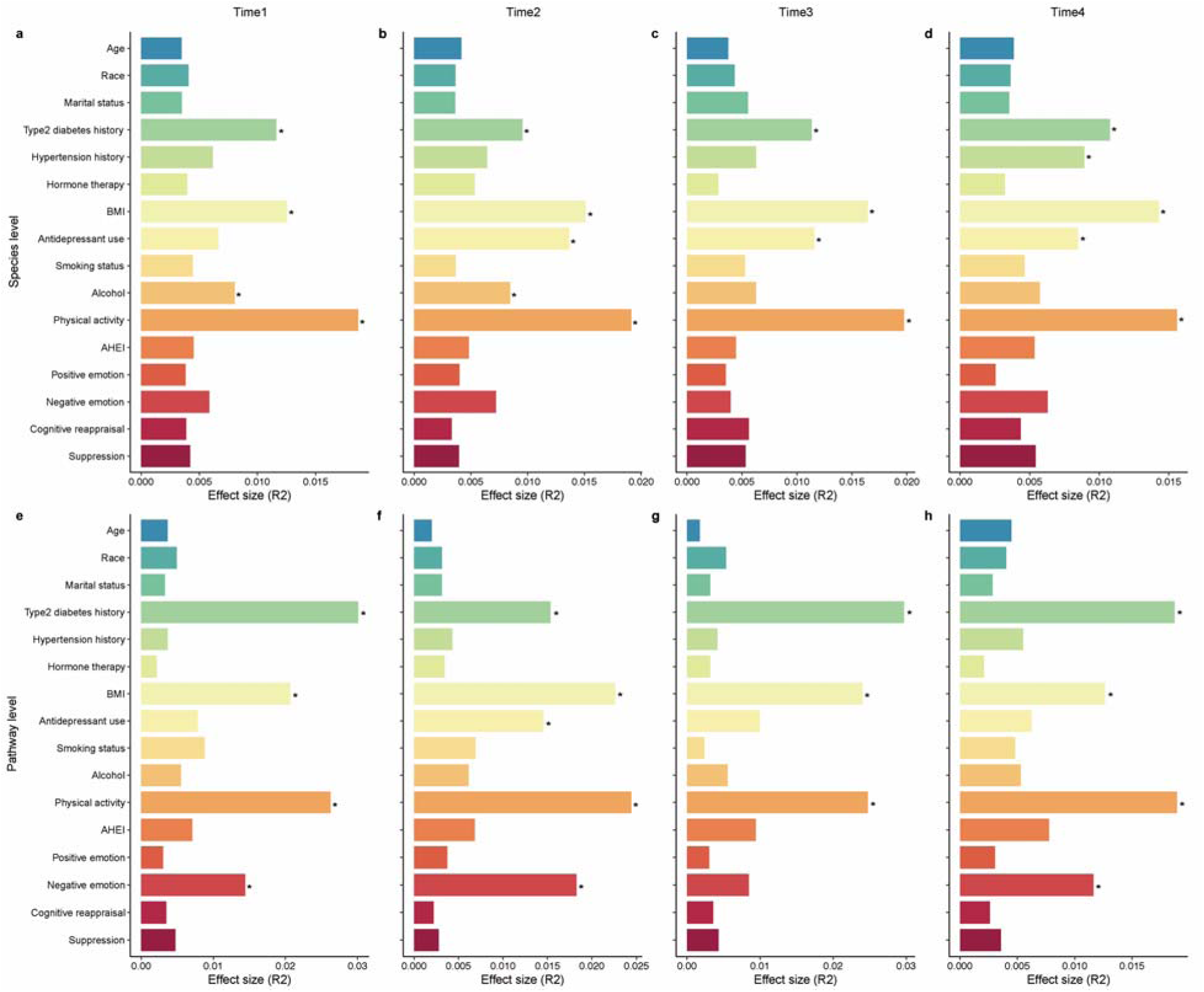
Gut microbiome associated host factors. The amount of variance (*r*^2^) explained by each host factor in the taxonomic (at the species-level, **a**-**d**) and functional (at the metabolic pathway-level, **e**-**h**) profiles determined by PERMANOVA analysis. All analyses were conducted based on all 787 metagenomes collected from 4 time points of 206 participants. The size of microbiome samples collected at first (**a, e**), second (**b, f**), third (**c, g**), and fourth (**d, h**) time points are 203, 197, 193, and 194. The asterisks denote significant associations (*p* **≤** 0.05). AHEI: alternate healthy eating index. The color of each bar represents a different host factor.

### Associations of emotion-related factors with species-level features and metabolic pathways of the gut microbiome

To explore the potential links of the gut microbial species and pathways with emotions and emotion regulation strategies, we performed per-feature testing using MaAsLin2 (Microbiome Multivariable Associations with Linear Models)^36^. These models account for within-individual association from the study’s repeated sampling design, as well as occasional missing observation of microbiome samples at some time points. Specifically, all models included each participant’s identifier as a random effect and simultaneously adjusted for significant potential confounders including physical activity, BMI, and type 2 diabetes history (i.e., the host factors identified as being significantly associated with variation in both taxonomic and functional profiles) as fixed effects. Top 10 <u>species-level features</u> from each emotion-related factor are summarized in Fig.4 and Table S2. Specifically, positive emotions were significantly and inversely associated with *Firmicutes bacterium CAG 94* and *Ruminococcaceae bacterium D16*, while negative emotions were significantly related to higher abundance of these same species (*q*≤0.25). Moreover, we found some species shared consistent relationships with emotions and emotion regulation strategies in the expected directions and with similar magnitude of association, although not all *q* values reached statistical significance. For example, *Bacteroides xylanisolvens, Firmicutes bacterium CAG 9*5, *Parabacteroides distasonis* were correlated with higher level of both positive emotions and cognitive reappraisal, as well as inversely correlated with both negative emotions and suppression. Further, we observed that positive emotions and cognitive reappraisal were associated with lower abundance of multiple species (i.e., *Anaeromassilibacillus sp An250, Bacteroides faecis, Blautia hydrogenotrophica, Clostridium bolteae CAG 59, Clostridium leptum, Firmicutes bacterium CAG 94, Ruminococcaceae bacterium D16, Sellimonas intestinalis, Streptococcus parasanguinis*), while negative emotions and suppression were correlated with higher abundance of these species.

**Fig. 4.**
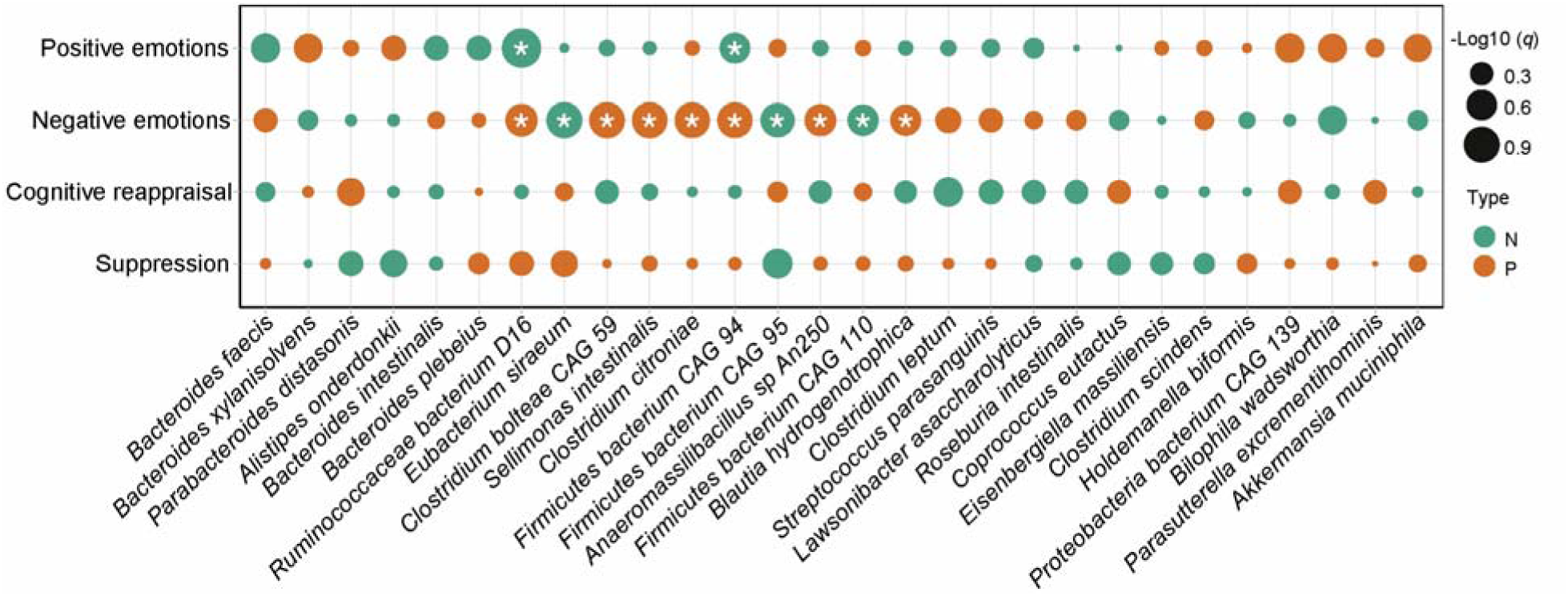
Associations of emotion-related factors with microbial species. Significant associations between positive and negative emotions and emotion regulation strategies and microbial species were identified using MaAsLin2 (Microbiome Multivariable Associations with Linear Models). Top 10 species-level features from each emotion-related factor are summarized (based on *q* value). Only statistically significant associations with *q* value **≤** 0.25 (Benjamini-Hochberg adjusted *P* value) are labeled with an asterisk. The size of each dot represents the – log10 (*q* value). The color of each dot represents the valence of the association: green=negative, orange =positive. All analyses were conducted based on all 787 metagenomes collected from 206 participants. See Table S2 for coefficients and exact p and q values on these microbial species.

We then investigated the emotion-related factors in relation to metabolic pathways, with the top 10 pathways from positive emotions, cognitive reappraisal, suppression, and negative emotions summarized in Fig.5 and Table S3. Positive emotions were significantly associated with higher abundance of three pathways (PWY0-1241: ADP-L-glycero-β-D-manno-heptose biosynthesis, PWY0-1298: superpathway of pyrimidine deoxyribonucleosides degradation, and FUCCAT-PWY: fucose degradation). Negative emotions were associated with a total of 63 metabolic pathways (Fig.5 and Table S4, *q*≤ 0.25). For example, negative emotion scores were significantly inversely correlated with pathways related to coenzyme A (CoA) biosynthesis (e.g., COA-PWY-1, PANTOSYN-PWY, COA-PWY, and PWY-4242), adenosine biosynthesis (e.g., PWY-6609, PWY-7219, PWY-7229, and PWY-6126), and pyrimidine deoxyribonucleosides salvage (PWY-7199). Suppression scores were negatively associated to pyrimidine deoxyribonucleotides de novo biosynthesis II (PWY-7187) and a pathway related to biosynthesis of propanoate (P108−PWY, Fig.5). However, we found no significant associations between cognitive reappraisal and any metabolic pathway.

**Fig. 5.**
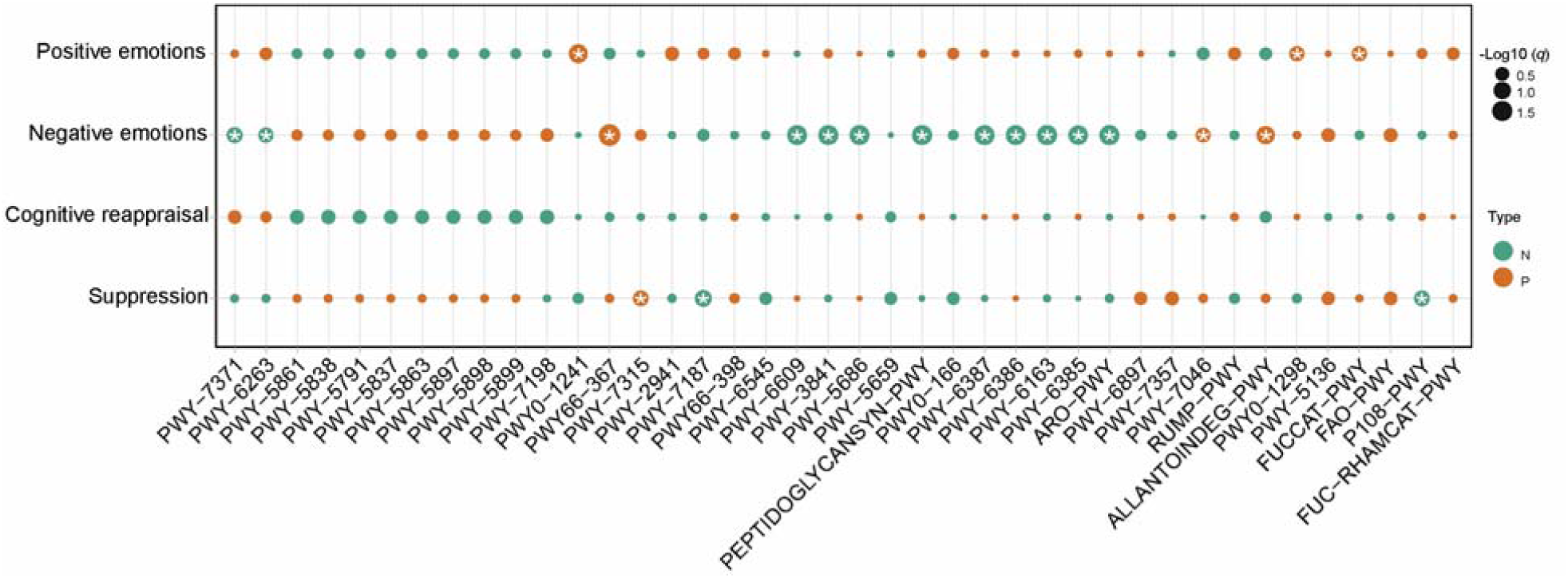
Associations of emotion-related factors with metabolic pathways. Significant associations of positive and negative emotions and emotion regulation strategies with metabolic pathways were identified using MaAsLin2. Top 10 metabolic pathways from positive emotions, negative emotions, cognitive reappraisal, and suppression were summarized. Only the statistically significant associations with *q* value **≤** 0.25 (Benjamini-Hochberg adjusted *P* value) are labeled with an asterisk. The size of each dot represents the –log10 (*q* value). The color of each dot represents the valence of the association: green=negative, orange =positive. All analyses were conducted based on all 787 metagenomes collected from 206 participants. See Table S3 for the annotations of these metabolic pathways.

## Discussion

In this discovery-based study, we examined associations between four emotion-related factors and the gut microbiome, leveraging a richly characterized cohort of 206 women. This study is the largest to date to evaluate associations of the gut microbiome with both positive and negative emotions and to assess the gut microbiome in relation to emotion regulation strategies. We found specific emotion-related factors were linked to microbiome diversity, as well as with certain species and metabolic pathways. These findings collectively offer early evidence suggesting emotions and emotion regulation strategies are related to the gut microbiome in ways that may have health-relevant implications.

A prior study identified the significant association of microbial diversity (i.e., Shannon diversity) with emotions among participants in the *Prevotella*-enterotype group. Moreover, both positive and negative emotions were associated with a novel genus (*PAC001043_g*) from the family Lachnospiraceae ^18^. Among the emotion-related factors included in our study, only suppression was significantly associated with alpha diversity. Notably, higher levels of suppression were associated with lower values on the Simpson diversity index and less Evenness in the gut microbial community at the species level. We also considered microbiome associations at the species and metabolic pathway levels. These analyses consistently demonstrated specific microbial associations of positive emotions and cognitive reappraisal in one direction and opposite associations with negative emotions and suppression. For example, lower levels of positive emotions and higher levels of negative emotions were associated with increased *Firmicutes bacterium CAG 94* and *Ruminococcaceae bacterium D16*. Interestingly, a multi-study, integrative analysis on 4,347 human stool metagenomes from 34 published studies including healthy and unhealthy individuals (e.g., having atherosclerotic cardiovascular disease, colorectal cancer, obesity) found *Ruminococcaceae bacterium D16* was less prevalent in healthy (vs. unhealthy) individuals^37^. Similarly, we observed that higher levels of positive emotions and cognitive reappraisal were inversely correlated with multiple other species (e.g., *Anaeromassilibacillus sp An250, Bacteroides faecis, Blautia hydrogenotrophica, Clostridium bolteae CAG 59, Clostridium leptum, Sellimonas intestinalis*, and *Streptococcus parasanguinis*), while higher levels of negative emotions and suppression were associated with increased level of these same species. Some of these species were previously reported as being associated with human disease related to both mental health and inflammatory disorders, conditions that have also been linked with emotional distress and dysregulation^38^. For example, *C. bolteae* has been found elevated in neuromyelitis optica spectrum disorders^39^, and *S. intestinalis* was found enriched in the gut microbiota of patients with schizophrenia^40^. *S. parasanguinis* is a dominant isolate of dental plaque and an opportunistic pathogen associated with subacute endocarditis^41^. Taken together, these findings suggest favorable emotional functioning, characterized by higher levels of positive emotions and lower levels of negative emotions, as well as better emotion regulation (i.e., greater use of cognitive reappraisal, and lower use of suppression), are associated with distinct compositional profiles of the gut microbiome at the species level. More research is needed to better understand the role of specific species in health, particularly for outcomes such as cardiovascular disease that have been related to emotions and emotion regulation.

When examining metabolic pathways, higher levels of negative emotions were associated with lower abundance of metabolic pathways in the biosynthesis of pantothenate and CoA. Pantothenate (i.e., vitamin B5) is the key precursor for the biosynthesis of CoA, a universal and essential cofactor involved in many metabolic reactions, including phospholipid synthesis, biosynthesis and degradation of fatty acids, and the tricarboxylic acid cycle^42^. Our results are in line with prior work that found moderate intake of pantothenic acid was related to lower odds of experiencing anxiety symptoms^43^. In addition, we found multiple metabolic pathways related to adenosine biosynthesis were significantly and inversely correlated with negative emotions. This is concordant with the fact that adenosine, one of the most ubiquitous and conserved neuromodulators in the central nervous system^44^, may have beneficial impacts upon depressive and anxious symptomatology^45-48^. However, how disruption of those metabolic pathways may interact with emotion-related factors warrants further investigation.

Our study has several limitations. First, this study was conducted among 206 adult women who are health professionals and mostly White. Therefore, our findings need to be validated by external studies of men, younger women, and larger or more diverse populations. Second, while bidirectionality in the association of emotion-related factors with the microbiome is likely, we cannot test for causality nor directionality in these relationships within this cross-sectional study. Our findings do identify rigorously assessed associations considering within-person correlation across microbiome variables due to the repeated sampling design as well as potential confounding factors, but future work using both human interventional studies and animal experiments is needed to ascertain directionality in these associations. These limitations are balanced by considerable strengths. Notably, the study includes the collection of multiple stool samples per participant, detailed phenotyping of the participants, and a validated measure of emotion regulation.

In summary, we found evidence that positive and negative emotions, as well as emotion regulation strategies, are related to specific aspects of the gut microbiome for both phylogenetically diverse organisms and specific metabolic pathways including pantothenate and CoA and adenosine. Together, these results connect emotional functioning and the human gut microbiome, highlighting the critical importance of incorporating the human microbiome in our understanding of emotion-related factors and their association with physical health.

## Methods

### Study population

Data are from a sub-study within the Nurses’ Health Study II (NHSII) cohort, an ongoing prospective cohort study of 116,429 US female registered nurses. At enrollment in 1989, all participants completed a questionnaire reporting on their demographics, lifestyle factors, and medical history. Biennial follow-up questionnaires have been mailed to all participants to update exposure information and disease diagnoses. Return of the completed questionnaires implied consent to use the data in ongoing research. The Mind-Body Study (MBS) was a NHSII sub-study of 233 postmenopausal women^21^. At study baseline in 2013, MBS participants (age: 49–67 years) signed a written informed consent form and completed a comprehensive online psychosocial assessment including measures of positive and negative emotions, as well as emotion regulation twice one year apart. The protocol asked participants to provide four stool samples to reduce measurement error—one pair collected 24–72 h apart and a second pair ∼6 months later. Women were eligible for inclusion in the current study if they completed at least one measure of emotion-related factors (i.e., positive emotions, negative emotions, or emotion regulation) and provided at least one valid stool sample (*N*=206). Among the 206 eligible participants, 179 (18, 8 and 1) participants provided 4 (3, 2 and 1) stool samples, respectively. This yielded 787 stool samples in total. The study protocol was approved by the institutional review boards of the Brigham and Women’s Hospital and Harvard T.H. Chan School of Public Health. Participants provided informed consent by returning questionnaires.

### Emotional Factors

#### Positive Emotions

Self-reported positive emotions were derived from 2 positively-worded items (“I was happy,” “I felt hopeful about the future”) from the 10-item Center for Epidemiological Studies Depression (CES-D-10) Scale^29^, which was completed at study baseline in 2013. Items were scored on a 4-point Likert scale with response options ranging from 1 (“Rarely or none of the time”) to 4 (“All of the time”). Items were averaged with higher scores indicating higher levels of positive emotions. A positive emotions score was assigned if participants completed both items (n=202). Internal consistency of the two items was acceptable (α = 0.75) and past work support a 2-factor structure for the CESD-10, with these two positively worded items loading on a positive affect factor^29,49^. We standardized the positive emotions score for analyses using the Z-score (Mean = 0, SD = 1).

#### Negative Emotions

We created an overall measure of negative emotions by pulling items from existing self-reported measures administered in MBS. Self-reported negative emotions were derived from combining data on 7 items: 4 items from the CESD-10^29^ (I was bothered by things that usually don’t bother me”, I felt depressed”, I felt fearful”, I was lonely”), and 3 non-redundant items from the Kessler Psychological Distress Scale, 6-item version^30^ (K-6; During the past month, about how often did you feel… “nervous”, “hopeless”, “restless or “fidgety”). The CESD-10 items were scored on a 4-point Likert scale with response options ranging from “Rarely or none of the time” to “All of the time”. The K-6 items were rated on a 5-point Likert with response options ranging from “None of the time to “All of the time”. To harmonize the K-6 items with the CESD-10 items, we combined two response options (i.e., “Some of time” and “A little of the time”) to create a 4-point Likert scale resulting in a similar set of response options. Items were averaged with higher scores indicating higher levels of negative emotions. A negative emotions score was created if participants completed 5 items (n=206). Internal consistency of the seven items was acceptable (α = 0.77). We standardized the negative emotions score for analyses using the Z-score (M = 0, SD = 1).

#### Emotion Regulation Strategies

Participants completed the Emotion Regulation Questionnaire, a validated 10-item scale that measures cognitive reappraisal (six items) and emotional suppression (four items)^31^. Items are scored on a seven-point Likert scale ranging from 1 (‘strongly disagree’) to 7 (‘strongly agree’). A total score is obtained for each subscale by averaging the items of the subscale, with higher scores reflecting higher use of each emotion regulation strategy. A cognitive reappraisal score was created if participants completed at least 5 items (n=204). For suppression, a score was created if participants completed at least 3 items (n=204). Internal consistency in this sample was good for the cognitive reappraisal subscale (α = 0.87) and acceptable for the suppression subscale (α = 0.76). Cognitive reappraisal and suppression scores were standardized for analyses (Mean = 0, SD = 1).

#### Sample collection, DNA extraction, and metagenome sequencing

All MBS participants were asked to provide 4 stool samples at multiple time points (Fig.1). Collection kits were mailed to participants with detailed instructions and return shipping to our biorepository via overnight courier within 24 hours of collection. Details on sample collection have been published elsewhere^21^. From 2013 to 2014, each participant provided up to two pairs of stool samples. Each pair of stool samples were collected from two bowel movements 24 to 72 h apart. The second set of two such samples were collected about six months after the first collection.

DNA purification from stool aliquots were performed according to standard protocols used in the Human Microbiome Project (HMP)^32,33^. Following previous work in MBS, construction and sequencing of sample libraries was conducted at the Broad Institute^50^. In particular, metagenome libraries were constructed using the Illumina TruSeq or Nextera method with ∼180nt inserts and sequenced on one of the Illumina HiSeq platforms (2500 or 4000) targeting a minimum of ∼2 Gnt/sample with 100nt paired-end reads.

#### Microbiome taxonomic and functional potential profiling

For the raw metagenomics sequencing data, low quality reads were discarded and reads belonging to the human genome were removed by mapping the data to the human reference genome with KneadData^35^. Microbial taxonomic profiling was performed using MetaPhlAn3^34^, which uses a library of clade-specific markers to provide panmicrobial (bacterial, archaeal, viral, and eukaryotic) profiling. We then performed functional profiling for metagenomes by applying HUMAnN3^34^, which maps DNA reads to a customized database of functionally annotated pan-genomes.

#### Covariates

Covariates included factors that can be related to both emotion-related factors and the gut microbiome; all were self-reported on the MBS questionnaire unless otherwise stated. Sociodemographics included age (continuous; based on date of birth queried at NHSII baseline in 1989), race/ethnicity (White, non-White; queried in 2005 on the NHSII biennial questionnaire), and marital status (married/cohabitating, never married/divorced/widow; queried in 2013 on the NHSII biennial questionnaire). Health-related factors (yes, no) included history of diabetes and hypertension (queried on every NHSII biennial questionnaire), hormone therapy use, and antidepressant use. BMI in kg/m2 was derived from height (queried in 1989) and weight^51^. Alcohol consumption in grams/day was derived from dietary information collected in 2013 using a validated semi-quantitative FFQ^52^. MET-hours per week was calculated from a validated questionnaire to quantify recreational physical activity based on 10 common activities (e.g., running, walking)^53^. Weekly physical activity was determined according to the number of minutes/week women reported spending in moderate to vigorous activity (<150 min/week, > 150 min/week; queried in 2013 on the NHSII biennial questionnaire)^53^.

Since the gut microbial composition and biosynthetic capacity are responsive to host diet and microbes in turn influence nutrients reaching the host through the metabolism of food^54^, we computed AHEI (the Alternate Healthy Eating Index 2010; a score that measures adherence to a diet pattern based on foods and nutrients most predictive of chronic disease risk) dietary score derived from the semiquantitative food frequency questionnaire (FFQ) data collected in 2013. The AHEI-2010 includes 11 variables: six components for which higher intakes are better (i.e., vegetables, fruit, whole grains, nuts and legumes, long-chain fats, and polyunsaturated fatty acids), one component for which moderate intake is better (i.e., alcohol), and four components that must be limited or avoided (i.e., sugar sweetened drinks and fruit juice, red/processed meat, trans fats, and sodium)^55^. All AHEI-2010 components were scored on a 0-to-10 scale, and then the total AHEI-2010 score was created by summing each component’s score. The AHEI-2010 score can range from 0 (non-adherence) to 110 (perfect adherence), with higher scores representing healthier diet patterns^56^.

#### Statistical analysis

We assessed Spearman correlations across emotion-related factors. Microbial diversity measures were calculated at the species and metabolic pathway levels, using the ‘vegan’ R (v.2.5-7) package. Measures of alpha diversity included Richness, Evenness, Simpson, and Shannon indices. For beta diversity for community composition and functional capacity over time, we used the Bray-Curtis dissimilarity and Principal Coordinates Analysis (PCoA). Differences in the gut microbiome over time were tested by the permutational multivariate analysis of variance (PERMANOVA) using the “adonis” function in R’s vegan package. We then applied PERMANOVA to estimate the effect size and significance of the effect of each covariate in relation to the gut microbial composition and metabolic pathway. All PERMANOVA tests were performed with 9999 permutations based on Bray–Curtis dissimilarity. To determine the statistical significance of associations of species or functional capacity with emotion-related factors while adjusting for covariates, MaAsLin2 (Microbiome Multivariable Associations with Linear Models, v.1.0.0) was used ^36^. Nominal *P* values across all associations were then adjusted for multiple comparisons using the Benjamini–Hochberg method with a target rate of 0.25 for *q* values^57^. All statistical analysis was performed with R (v.3.6.3) and SAS (v.9.4, 2013).

## Data availability

The data that support the findings of our study are available from Brigham and Women’s Hospital and Harvard T.H. Chan School of Public Health. Restrictions apply to the availability of these data, which were used under license for our study. Data are available (https://sites.google.com/channing.harvard.edu/cohortdocs/) with the permission of Brigham and Women’s Hospital and Harvard T.H. Chan School of Public Health.

## Author contributions

S.S.T. and L.D.K. designed and implemented the Mind Body Study within the Nurses’ Health Study II. L.D.K. and Y.-Y.L. conceived and designed the project. S.K. and A.J.G served as lead for conceptualization, performed the data analysis, and wrote the manuscript. L.D.K., Y.-Y.L., S.S.T., T.H., and A.T.C. edited the manuscript.

## Acknowledgements

We thank Jacqueline Starr, Scott Weiss, Susan Korrick, Qi Sun, Xu-Wen Wang, Zheng Sun, Andrea Aparicio, and Tong Wang for valuable discussions. We thank the participants and the staff of the Nurses’ Health Study II for their valuable contributions.

